# Inferring intrinsic population growth rates and *per capita* interactions from ecological time-series

**DOI:** 10.1101/2024.05.07.592896

**Authors:** Phuong L. Nguyen, Francesco Pomati, Rudolf P. Rohr

**Affiliations:** Department of Biology, University of Fribourg, Chemin du Musée 10, CH-1700 Fribourg, Switzerland; Department of Aquatic Ecology, Swiss Federal Institute of Aquatic Science and Technology (Eawag), Überlandstrasse 133, CH-8600 Dübendorf, Switzerland

**Keywords:** Lotka Volterra map, intrinsic growth rate, *per capita* interaction strength, time-series data, weighted multivariate regression

## Abstract

Knowledge about the *per capita* interactions between organisms and their intrinsic growth rates, and how these vary over environmental gradients, allows understanding and predicting species coexistence and community dynamics. Estimating these crucial ecological parameters requires tedious experimental work, with isolation of organisms from their natural context. Here, we provide a novel approach for inferring these key parameters from time-series data by using weighted multivariate regression on the *per capita* growth rates of populations. Beyond the validation of our approach on synthetic data, we reveal from experimental data an expected allocative trade-off between grazing resistance and rapid growth in algae. Application of observational data suggests facilitation between cyanobacteria and chrysophyte, indicating a possible explanation for cyanobacteria bloom. Our approach offers a way forward for inferring *per capita* interactions and intrinsic growth rates directly from natural communities, providing realism, mechanistic understanding of eco-evolutionary dynamics, and key parameters to develop predictive models.

## Introduction

To adapt to our rapidly changing planet, ecologists must be able to understand and predict the responses of entire ecosystems. This requires the study of species not as individuals but as interacting agents who collectively determine the emergent properties of complex and dynamic communities (Bascompte & Jordano, 2007; Cohen *et al*., 2009; Godoy *et al*., 2018). As such, most modern ecological and evolutionary theoretical models are founded on two key parameters (Chesson, 2000; Vincent & Brown, 2005; HilleRisLambers *et al*., 2012; Saavedra *et al*., 2017) that are essential for understanding and modelling how organisms interact (Turchin, 1999): the intrinsic growth rate of a population and the *per capita* interaction coefficient. By definition, these factors quantify, respectively, the *per capita* rate of change of a population at a low density, meaning in the absence of any limitations, and the effect that co-occurring organisms have on each other’s abundance. These key parameters are essential for understanding and predicting species coexistence, community composition, and ecosystem biodiversity (Chesson, 2000; Vincent & Brown, 2005; Bascompte & Jordano, 2007; HilleRisLambers *et al*., 2012; Baert *et al*., 2016; Saavedra *et al*., 2017; Bartomeus *et al*., 2021). However, direct measurements or more practical estimations of these parameters remain challenging for ecologists.

This challenge currently stems from the complexity of the experimental setups that are required to measure these parameters and variations in how interactions are measured due to differences in dimensions and units (Berlow *et al*., 2004; Arditi *et al*., 2021). As an example dating back to 1969, Vandermeer estimated all pairwise *per capita* interactions and intrinsic growth rates of four species of protozoa by fitting experimental data of monocultures and bi-cultures using the Lotka-Volterra multi-species model (Vandermeer, 1969), which has served as the foundation of most theoretical models in ecology and evolution (Vandermeer, 1969; Turchin, 1999; Chesson, 2000; HilleRisLambers *et al*., 2012; Saavedra *et al*., 2017). This work required at least 10 time-series (four monocultures and six bicultures), without replication. Furthermore, the *per capita* interaction is not the only measurement of interaction. Subsequent work by Laska & Wootton (1998) identified three additional representations of the concept of interaction, including: (i) the Paine’s index that reflects the difference in abundance of a community containing all species and lacking a focal species; (ii) the Jacobian matrix that shows the direct effect of one species on the total abundance of another species; and (iii) the inverted Jacobian matrix that includes both direct and indirect effects, such as apparent competition and competition via resources (Bender *et al*., 1984; Berlow *et al*., 2004). It is worth noting that these three concepts of interaction either require populations at ecological equilibrium, such as Paine’s index, or are density-dependent, such as the Jacobian matrix. However, to understand community dynamics in terms of governing mechanisms and to develop realistic mechanistic models, it requires the intrinsic growth rate and the *per capita* interactions inferred in Vandermeer’s work and in later experimental studies (Levine & HilleRisLambers, 2009; Bartomeus *et al*., 2021; Van Dyke *et al*., 2022).

To overcome labour-intensive experimental work, Sugihara and collaborators use a weighted multivariate multilinear regression, the S-map method, to infer the Jacobian matrix directly from observational data (Sugihara, 1994; Deyle *et al*., 2016). While this technique does not require populations to be at ecological equilibrium, as one can study the temporal change of the Jacobian elements, it still infers elements of the Jacobian matrix and not the *per capita* interaction strengths (Berlow *et al*., 2004; Chang *et al*., 2021; Arditi *et al*., 2021). Moreover, the intraspecific components of the Jacobian matrix, determined by the diagonal elements, are often neglected because they entangle intrinsic growth rates and *per capita* intraspecific interactions. Regardless, an inference of both intrinsic growth rate and *per capita* interaction strength, as was done originally in Vandermeer (1969), is required for studying community coexistence (Chesson, 2000; Saavedra *et al*., 2017), productivity (Parain *et al*., 2019), and mechanisms underlying community and evolutionary dynamics, such as allocative trade-offs between intrinsic growth and *per capita* interaction strengths (Vincent & Brown, 2005; HilleRisLambers *et al*., 2012).

Thus, to directly infer the intrinsic growth rate and *per capita* interaction strength from complex, dynamic communities, we herein propose a novel approach called Lotka-Volterra map (LV-map), which combines the strength of the S-map’s inference ability and the mechanistic understanding of population’s ecological nature in the Lotka-Volterra model (Lotka, 1925; Volterra, 1931). The key innovation of our approach is to estimate, from observational data, the intrinsic growth rate and *per capita* interaction parameters, and their potential variation with time and environmental conditions, that are key for understanding and modelling ecological communities. To do so, we use the *per capita* growth rate as the response variable for weighted multivariate multilinear regression. In this way, the intercept and the slope of this regression naturally correspond to the intrinsic growth rate and *per capita* interaction strength. LV-map is not simply a multivariate regression because parameter inference is performed at each time point of the time-series, which enables the detection of potential time variations in these parameters. We first demonstrate on synthetic data that our approach provides the desired and correct parameters, then we illustrate its success on empirical data from both controlled experimental communities as well as observations. Subsequently, we explain the key differences between the Jacobian elements inferred by the S-map method, and the *per capita* interaction estimated from our LV-map. Our approach therefore serves as a robust tool for addressing ecological and evolutionary questions both within experimental setups and in natural communities.

## The Lotka-Volterra map approach

### Mechanistic basis of the approach

To build up the LV-map, it is essential to realise that population dynamics are governed by the birth and death of individual organisms. A key metric for monitoring changes in population sizes is naturally the *per capita* rate of change, which is the difference between the *per capita* birth and death rates. These *per capita* rates, in turn, are functions of the population densities, that is, the so-called density dependence of population growth (Turchin, 1999).

From a mathematical standpoint, in a community of *S* populations (which could be at the species, phenotypic, or genotypic levels), the changes in population densities are represented by their *per capita* rates, which are given by the log-ratio of population density changes: ln(*n*_*i*_(*t*+1)*/n*_*i*_(*t*)) = *λ*_*i*_(**n**(*t*), **e**(*t*)) (Turchin, 1999; Vincent & Brown, 2005). This *per capita* rate depends on all biotic and abiotic factors, represented respectively by the population densities **n**(*t*) and environmental conditions **e**(*t*). We can now incorporate our two key terms governing the *per capita* rate of change: the intrinsic growth rate and the limits imposed by interactions within and between species (Turchin, 1999; Sibly & Hone, 2002; Vincent & Brown, 2005). The former represents the intrinsic growth of a population in the absence of limitations, represented as the *per capita* rate of change when population densities are extremely low, that is, *r*_*i*_(*t*) = *λ*(**0, e**(*t*)). The latter refers to the regulation by both inter- and intraspecific *per capita* interactions, which is represented by the partial derivative of the *per capita* rates of change, *α*_*ij*_(*t*) = *∂λ*_*i*_(**n**(*t*), **e**(*t*))*/∂n*_*j*_(*t*). With population densities recorded in time-series, for each time point, we can approximate the *per capita* rates of change by a multivariate function of these population densities as follows:

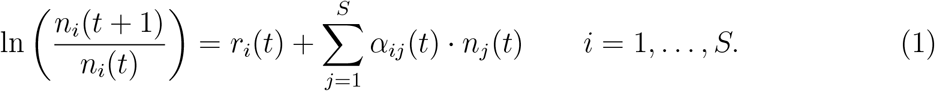

In this approximation, the intercepts correspond to the intrinsic growth rates, while the slopes represent the *per capita* interaction strengths (Vandermeer, 1969; Berlow *et al*., 2004; Vincent & Brown, 2005; Arditi *et al*., 2021). Equation 1 is, in fact, similar to the multi-species Lotka-Volterra model, with one subtle but fundamental difference—we do not assume constant values for *r*_*i*_(*t*) and *α*_*ij*_(*t*). This requires a weighting parameter *θ* that determines how *r*_*i*_(*t*) and *α*_*ij*_(*t*) vary with time. Details of *θ* are further explained in the next section.

### Weighted multilinear multivariate regression

We now show that by using the *per capita* growth rate of the population *λ*_*i*_(**n**(*t*), **e**(*t*)) as the response variable, the LV-map is, in fact, a multivariate non-linear autoregressive process. Equation 1 can be rewritten as

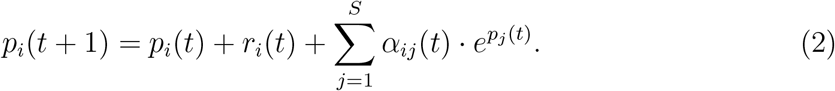

where *p*_*i*_(*t*) = ln *n*_*i*_(*t*). The parameters *r*_*i*_(*t*) and *α*_*ij*_(*t*) can be inferred by maximum likelihood techniques, which is equivalent to minimizing the residual sum of squares (RSS):

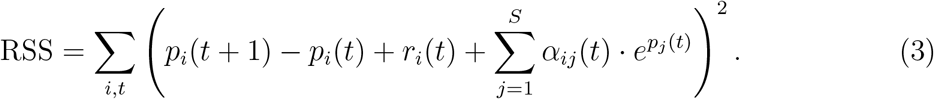

Putting back *n*_*i*_(*t*) in the above equation, we obtain

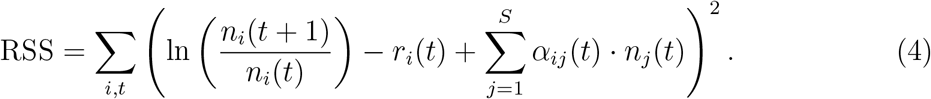

This is the exact same expression for a multivariate regression with *Y*_*i*_(*t* + 1) = ln(*n*_*i*_(*t* + 1)*/n*_*i*_(*t*)) the response variables and *n*_*i*_(*t*) the explanatory variables.

As the values of *r*_*i*_(*t*) and *α*_*ij*_(*t*) are free to vary with different conditions, and thus with time, we have to introduce a weighting kernel which weights each variable relative to the time point *t* that we consider. The state-space weighting kernel, introduced in the S-map (Sugihara, 1994; Deyle *et al*., 2016), takes into account the difference between a variable at time point *t* of focus and variables at other the time point *l* is defined as

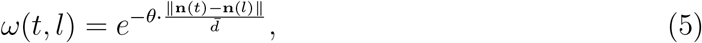

where ∥**n**(*t*) *−* **n**(*l*)∥ is the Euclidean distance between the vector of population densities **n**(*t*) at time point *t* and the one at time point *l*, and 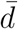 is the average Euclidean distance computed across all time-points *l*. The parameter *θ* determines the strength of the weighting kernel. This allows to define 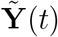 and 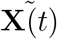 respectively as the matrix of weighted response and explanatory variables. Given that

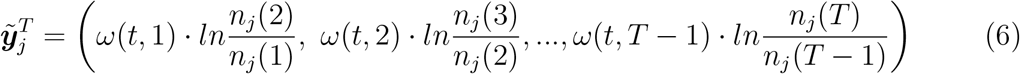

the column vector *j* of matrix 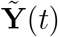, and

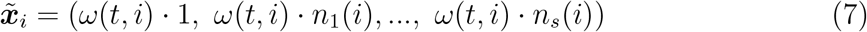

the row vector *i* of matrix 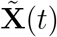. Here, 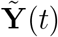 is of size (*T −* 1) *× S* and 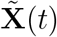 is of size (*T −* 1) *×* (*S* + 1), where *T* is the total data points of the time-series.

From these two matrices, we compute the lease square estimation of all parameters at time *t* by matrix computation:

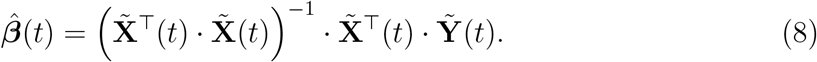

The matrix 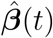 contains the estimations of the intrinsic growth rates of all populations as well as all *per capita* interaction strength. For instance, its column *j* is given by the column vector 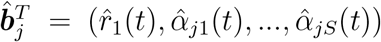 and provides the estimations of the intrinsic growth rate of the population *j* and the interaction of other populations in the communities on the population *j*. Note that 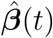 is a matrix of size (*S* + 1) *× S*. The mathematical proof can be found in chapter 3.2 of Hastie *et al*. (2001) and chapter 3.3 of Mardia *et al*. (1979). The strength of the weighting kernel, *θ*, is determined by cross validation (Supporting Information). Finally, we provided an estimation of the standard error for each 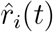 and 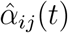 (Supporting Information).

### Fundamental differences between the LV-map and the S-map

Comparing the LV-map and S-map is essential as both methods use weighted multivariate multilinear regression, which has been widely discussed in time-series analysis and standard statistics (Hastie *et al*., 2001; Holger Kantz, 2004), although with different response variables. Specifically, with LV-maps, we model the *per capita* birth and death processes by considering the *per capita* growth rate, while the S-map assumes no process, and models directly the total growth rate. It is this subtlety that set a critical difference between these two methods, shifting the purpose from prediction to explanation by rendering the LV-map the capacity to delve deeply into mechanisms underlying the eco-evolutionary properties of communities. While the LV-map is able to infer the intrinsic growth rate and the *per capita* interaction strength, the S-map estimates the Jacobian elements by inferring parameters of the following model:

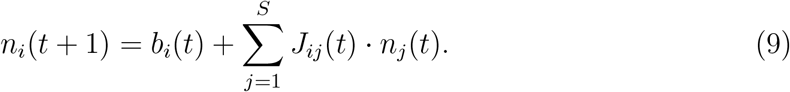

In the S-map, the intercept *b*_*i*_ carries no biological meaning, and the Jacobian elements can be related to *r*_*i*_(*t*) and *α*_*ij*_(*t*) of the LV-map as follows:

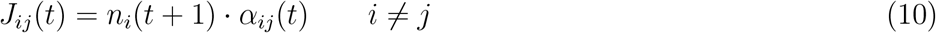

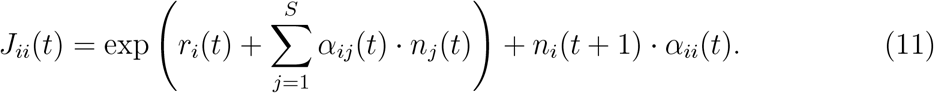

These Jacobian elements of S-maps indicate the total effect of one population on the growth of another population, i.e. the effect includes population densities as expressed in Equations 10 and 11. The off-diagonal Jacobian elements (*J*_*ij*_(*t*) for *i≠ j*, Equation 10 represent the *per capita* interaction strengths multiplied by the population densities, while the diagonal elements (*J*_*ii*_(*t*), Equation 11 include both the *per capita* intraspecific interaction (*α*_*ii*_(*t*)) and the intrinsic growth rates (*r*_*i*_(*t*)). Overall, while these methods are relatable, the convoluted terms generated by the Jacobian elements in Equations 10 and 11 make it extremely challenging to pull out values for the intrinsic growth rate and *per capita* interaction strength, especially as compared to the direct estimation of these values from the LV-map, as represented by Equation 1.

## Results

### Validating the LV-map using synthetic data

As our first application of the LV-map, we estimated the *r*_*i*_(*t*) and *α*_*ij*_(*t*) parameters from a discrete-time Lotka-Volterra model with environmental noise (Supporting Information). Our inferred parameters matched the true values, which was expected as this data is simulated from the Lotka-Volterra model (Figure 1). Note that the population dynamics of the simulated data demonstrate a cyclic behaviour, but the parameters (*r*_*i*_ and *α*_*ij*_) used for the simulation are constant (Figure 1a - c). Here, the key difference between our LV-map and the S-map is that the net interactions (Jacobian elements) inferred by the S-map change with respect to population density. As by definition, these are the total effects of one population on the others, and hence, are naturally density-dependent, as shown in Equation 10 (Figure 1b and d). This difference is also shown in the cross validation results, where *θ* = 0 with LV-map, and *θ >* 0 with S-map (Figure 2). When comparing to the LV-map, the net interaction of population 3 on population 2 inferred by the S-map is stronger due to the high density of the population 2, but not because the *per capita* interaction strength itself is inherently stronger (Figure 1c, d). More importantly, we could use LV-map to infer both intrinsic growth rates and intraspecific interactions, and were thus not limited to interspecific interactions as in previous studies (Paine, 1992; Deyle *et al*., 2016; Chang *et al*., 2021).

**FIGURE 1.**
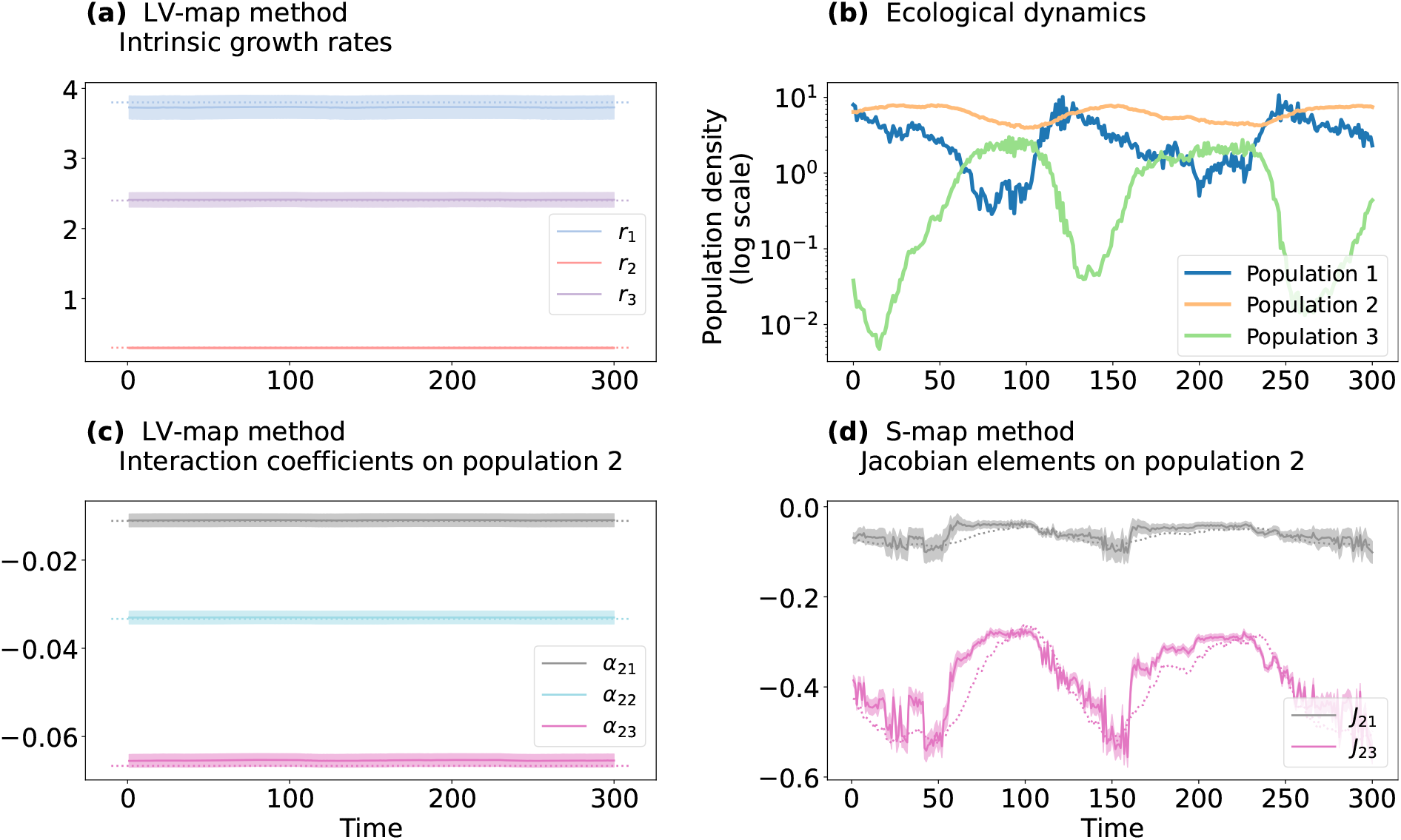
Estimations of the synthetic cyclic system, comparing the LV-map to the S-map. (a) Intrinsic growth rates estimated from LV-map. (b) Population dynamics of a Lotka-Volterra system with three competing populations. (c) *Per capita* interaction strengths of all populations on population 2, as estimated from LV-map. (d) Off-diagonal Jacobian elements for population 2, as estimated by S-map. Solid lines represent estimated values, and shaded areas illustrate the standard error. Dotted lines represent true values. Parameter inference for populations 1 and 3 and cross validation results can be found in Figure S1. Parameters used for the simulations are **r** = [3.8; 0.3; 2.4], ***α*** = *−*[(0.2, 0.4, 0.8); (1*/*90, 1*/*30, 1*/*15); (0.2, 0.2, 0.6)]. Environmental noise follows a normal distribution 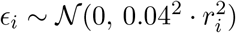

**FIGURE 2.**
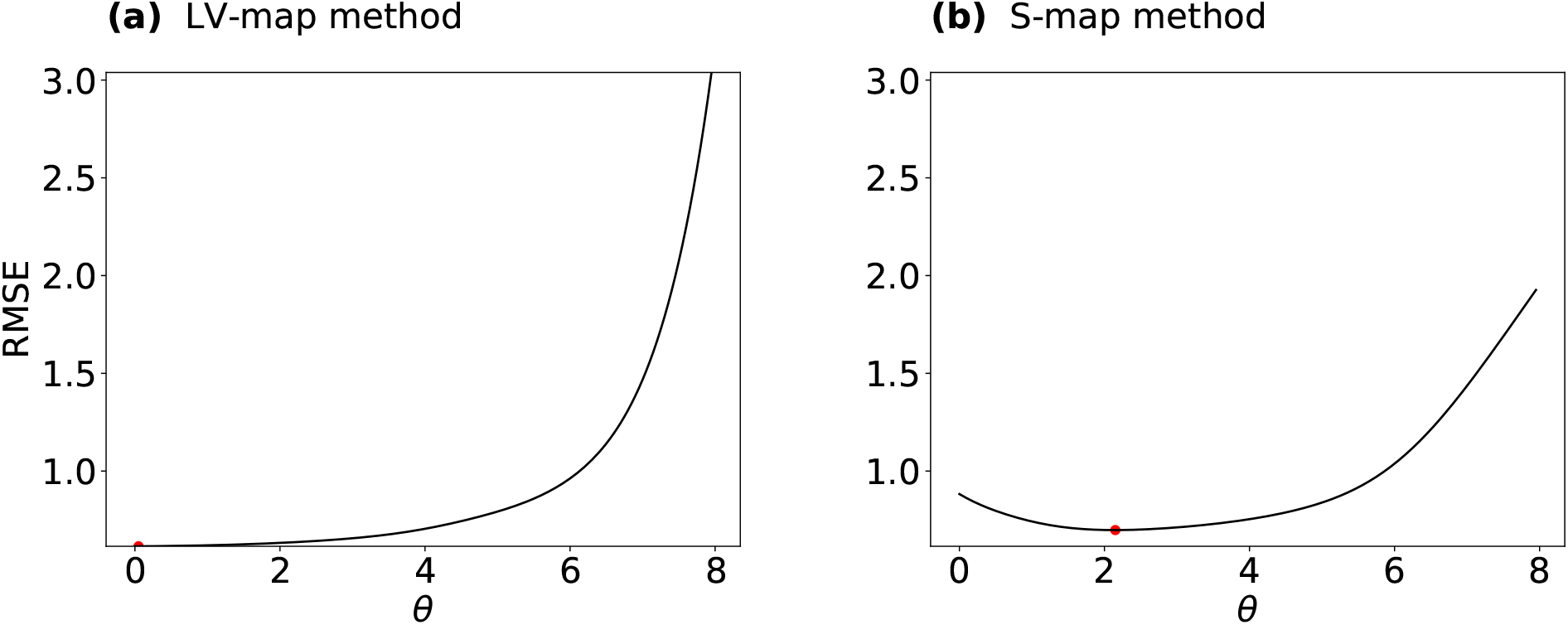
Cross validation results for the cyclic Lotka-Volterra model. (a), LV-map method. (b), S-map method. Red points indicate the value of *θ* with the smallest value of RMSE.

Then we test the LV-map against chaotic dynamics, which often is the case in nature. Figure 3a and c shows that while the LV-map can accurately infer the *r*_*i*_(*t*) and *α*_*ij*_(*t*), Figure 3d shows that in contrast, the S-map seems unable to infer the correct Jacobian elements. This is probably due to abrupt changes in population densities in the chaotic Lotka-Volterra dynamics, which challenges the inference of the density-dependent Jacobian elements. These two applications to synthetic data prove that we can effectively retrieve the expected values of the key ecological parameters governing community dynamics, with varying levels of complexity.

**FIGURE 3.**
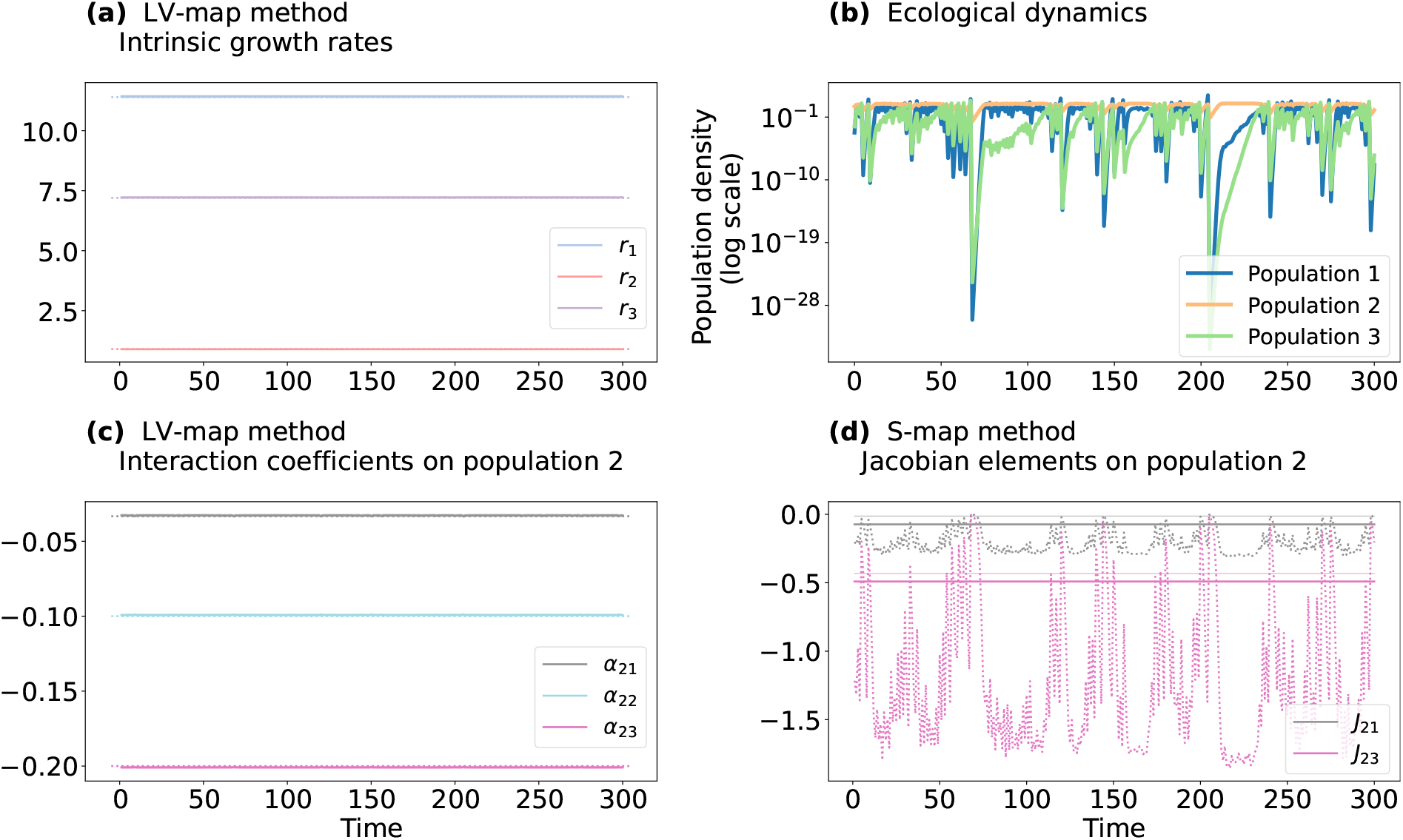
Estimations of a synthetic chaotic system, comparing the LV-map to the S-map. (a) Intrinsic growth rates estimated from LV-map. (b) Population dynamics of a Lotka-Volterra system with three competing populations. (c) *Per capita* interaction strengths of all populations on population 2, as estimated from LV-map. (d), Off-diagonal Jacobian elements for population 2, as estimated by S-map. Solid lines represent estimated values, and shaded areas illustrate the standard error (here, the standard errors are too small that they are almost not visible on the graphs). Dotted lines represent true values. Estimation of parameters for populations 1 and 3 and cross validation results can be found in Figure S2. Parameters used for the simulations are **r** = [11.4; 0.9; 7.2], ***α*** = *−*[(0.6, 1.2, 2.4); (0.033, 0.1, 0.2); (0.6, 0.6, 1.8)]. Environmental noise follows a normal distribution 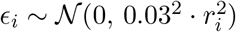.

### Analysing real-world data using LV-map

To then perform a proof-of-concept test of our approach using empirical data, we first applied the LV-map to a phytoplanktonic predator-prey system from Blasius *et al*. (2020) and Yoshida *et al*. (2003) (Supporting Information). We then apply it to a high-frequency time-series of five phytoplankton groups from lake data (Supporting Information).

The first set of phytoplanktonic experimental data was obtained from chemostat experiments wherein rotifers (*Brachionus calyciflorus*) and algae (*Monoraphidium minutum* inBlasius *et al*. (2020) and *Chlorella vulgaris* in Yoshida *et al*. (2003)) were cultivated together in constant environmental conditions with daily density measurements (Figure 4a - c and Figure S5a - c). These measurements were conducted on the clonal level for algae, resulting in three time-series data sets for each system.

**FIGURE 4.**
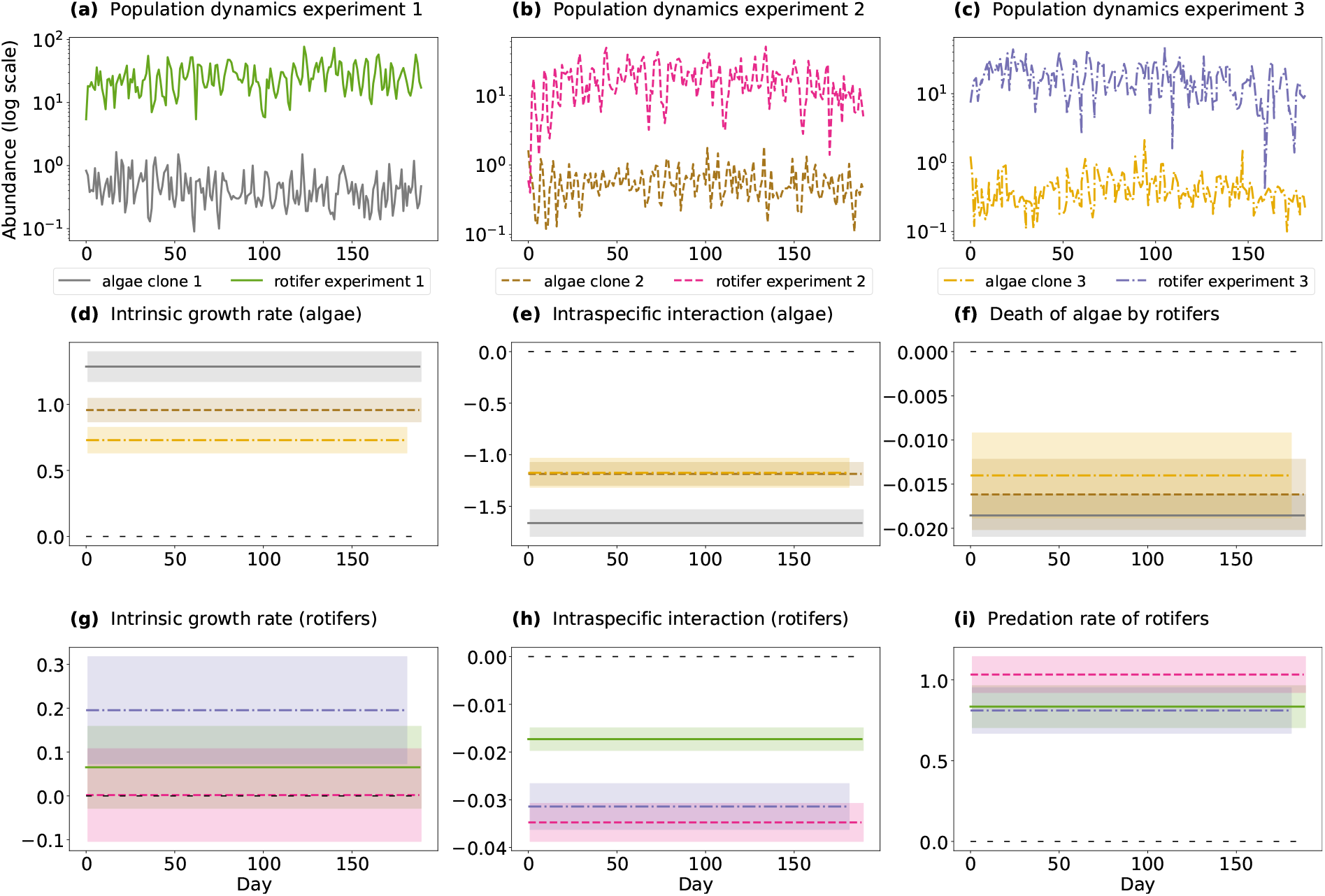
LV-map estimation of time-series data of the interactions between algae and rotifers from Blasius et al.Blasius *et al*. (2020). (a - c), Population dynamics. (d), Intrinsic growth rate of algae. (e), Intraspecific interactions between algae. (f), Effect of rotifers on algae. (g), Intrinsic growth rate of rotifers. (h), Intraspecific interaction between rotifers. (i), Effect of algae on rotifers. Shaded areas represent the standard error.

We thus applied the LV-map on all six time-series data and inferred the intrinsic growth rates and *per capita* interaction rates. The inferred intrinsic growth rates of all algae clones are positive, suggesting their autotrophic nature (Figure 4d and Figure S5d). The intraspecific interactions are negative, though, suggesting that the algae compete for nutrients (Figure 4e and Figure S5e), and the negative effect of rotifers on algae indicates that the algae are eaten by rotifers (Figure 4f and Figure S5f). Interestingly, in the experiment of Blasius *et al*. (2020) with algal clone 2, the intrinsic growth rate of the rotifers is almost zero, indicating that this predator cannot survive without the algae (Figure 4g). However, in the experiments with the other algal clones, the rotifers have slightly positive intrinsic growth rates, implying that they may exhibit some form of mixotrophic behaviour (Figure 4g, and Figure S5g). This result could also be explained by the ability of rotifers to exploit other resources in the system, such as particles or dissolved organic carbon sources. Overall, the effect of algae on rotifers is positive, suggesting that rotifers thrive on algae (Figure 4h and Figure S5h), though the negative interactions between rotifers indicate that they compete with each other (Figure 4i and Figure S5i).

The different parameters inferred by LV-map for the different algal clones suggests an allocative trade-off between reproduction and defensive traits. Allocative trade-off is common in nature as organisms always experience limited resources which are spent on growth, reproduction, defence, and so on (Strauss *et al*., 2002; Cotter *et al*., 2004; Yoshida *et al*., 2004). Thus, the more energy invested in one trait, the lesser energy is left for the others. Our results from Blasius’ experiment clearly show that the clone with the highest intrinsic growth rate (better reproduction) exhibits the strongest intraspecific competition and is also the most strongly grazed upon (less defence), and vice versa for the clone with the lowest intrinsic growth rate (Figure 4d - f). The experiments of Yoshida *et al*. (2003) exhibit an analogous pattern, despite some fluctuations in parameter values (Figure S5).

Another important result from LV-map is the value of the weighting parameter *θ* from cross validation. The best value for *θ*, which determines how the *r*_*i*_ and *α*_*ij*_ vary with time, was zero in four out of the six total data sets with *θ* = 0 for all experiments in Blasius *et al*. (2020)(Figure 5). This suggests that overall, the experimental predator-prey dynamics reflects the original Lotka-Volterra model, i.e., the parameters are time independent in a controlled experimental environment. In two other experiments in Yoshida *et al*. (2003), we have *θ≠* 0, suggesting a deviation from the original Lotka-Volterra model even though all experiments follow the same procedure (Figure S4). This could be because these experiments were short (20 days compared to 190 days in Blasius *et al*. (2020)), indicating that this length might be insufficient for the algorithm to fully capture the dynamics. Moreover, in one of these two experiments (Figure S4b), *θ* is, in fact, close to zero, implying that this dynamic could indeed follow the Lotka-Volterra model.

**FIGURE 5.**
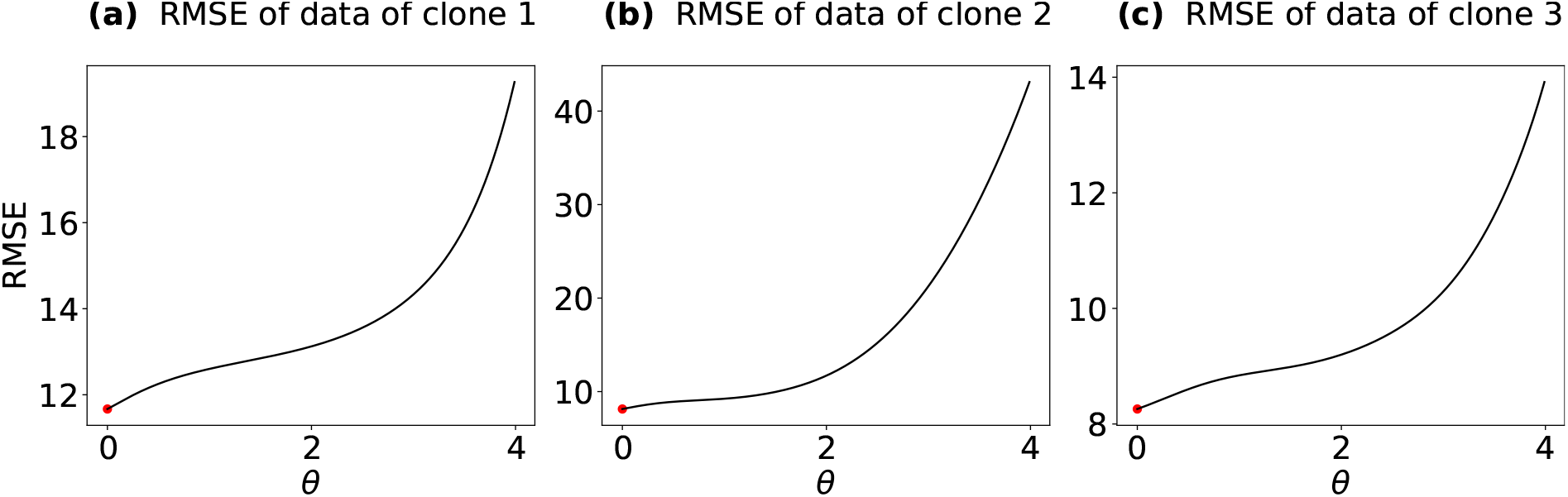
Cross validation results for experimental data of Blasius et al Blasius *et al*. (2020). Results from experiment 1 (a), experiment 2 (b), and experiment 3 (c). Red points indicate the value of *θ* with the smallest value of RMSE.

Overall, the application of the LV-map on experimental data shows that we can effectively estimate the intrinsic growth rates and *per capita* interactions in controlled experimental time series, avoiding the need for isolating species in monoculture, biculture, or other artificial setups as in previous studies (Vandermeer, 1969; Levine & HilleRisLambers, 2009; Bartomeus *et al*., 2021; Van Dyke *et al*., 2022). In addition, these estimated parameters allow elucidating eco-evolutionary interactions and can be directly implemented in LV-type models of community dynamics.

As a second real-world application of LV-map that tests its function on comparatively noisy data, we used high-frequency (daily) time-series data from Lake Greifensee, Switzerland. The data were collected by automated underwater imaging between March 2019 and December 2022 (Merz *et al*., 2021) (Supporting Information, Figure S6). We applied the LV-map to five phytoplankton groups, namely cyanobacteria, green algae, chrysophytes, diatoms, and cryptophytes (Figure 6). Here, our inferred intrinsic growth rates for phytoplankton are generally positive, except for cyanobacteria, which displayed an intrinsic growth rate of nearly zero (Figure 6 a). This suggests that cyanobacteria grow quite slow in nature, compared to other algae. Most of the inferred inter- and intraspecific interactions are small, except for chrysophytes, which had a large effect on the other groups (Figure 6 b-f). In some cases, we observed positive interspecific interactions, suggesting facilitating effects between groups (Figure 6 b,d). In particular, the positive interaction between the chrysophytes and the cyanobacteria may be an important hypothesis to test in follow-up studies in relation to cyanobacterial blooms — these events are becoming more and more common worldwide, they are difficult to predict and generally explained as a function of only abiotic environmental drivers (Huisman *et al*., 2018). In addition, these parameters show temporal fluctuations that can be reconciled with a seasonal variation of the environmental context, as would be expected for a dynamic system (Figure S7). Overall, this second application shows that we can effectively estimate the intrinsic growth rates and *per capita* interactions in natural phytoplankton communities using noisy *in situ* data, and retrieve valuable information to understand and model real-world problems (e.g. cyanobacterial blooms).

**FIGURE 6.**
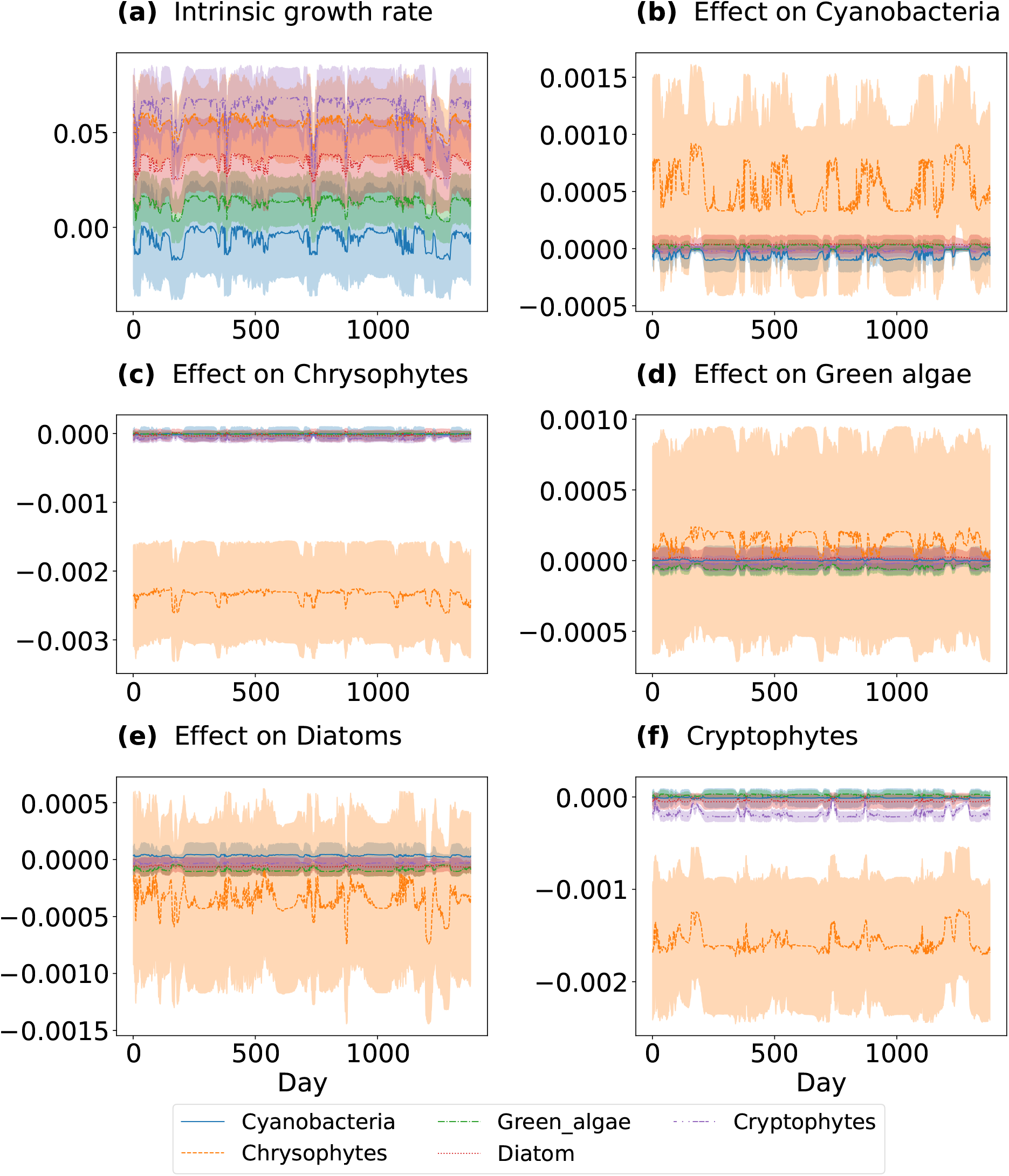
LV-map parameter inference from time-series data of five autotrophic groups in Lake Greifensee, Switzerland. (a), Intrinsic growth rates. (b), Effect of other groups on cyanobacteria. (c), Effect of other groups on green algae. (d), Effect of other groups on chrysophytes. (e), Effect of other groups on diatoms. (f), Effect of other groups on cryptophytes. The shaded area represents the standard error.

## Discussions

In this article, we present a new approach, termed LV-map, which we validated on synthetic, experimental and observational data. The LV-map is a weighted multivariate regression that offers a robust method for inferring ecological parameters without isolating organisms from a community context — a key issue in studies on community dynamics. These key ecological parameters, namely, *per capita* interaction and intrinsic growth rate, dictate the eco-evolutionary dynamics of communities. Together, they determine possibilities for species coexistence (Chesson, 2000; Saavedra *et al*., 2017), structures of communities in nature (Bascompte & Jordano, 2007; Cohen *et al*., 2009), and the relationships between biodiversity and ecosystem functioning (Baert *et al*., 2016, 2018; Bartomeus *et al*., 2021).

The LV-map approach presents subtle differences with the S-map that expand its utility in ecology and evolution: LV-map can infer the *per capita* interactions instead of the elements of Jacobian matrix by using the *per capita* rate of population change rather than the total densities. Consequently, the intercepts and the slopes of the LV-map approach correspond naturally to the intrinsic growth rates *r*_*i*_(*t*) and *per capita* interaction strengths *α*_*ij*_(*t*), respectively. The determination of these two values often stymies studies into eco-evolutionary outcomes. As the multivariate regression is weighted, estimations made using LV-map can also be made time-dependent, such that it has the capacity to detect variations in *r*_*i*_(*t*) and *α*_*ij*_(*t*) across time.

One of the highlights of the LV-map is that it presents an alternative to the laborious experimental work that is normally required to determine *r*_*i*_(*t*) and *α*_*ij*_(*t*), as it infers these values directly from time-series data and thus offers the opportunity to study these parameters in natural communities instead of experimental ones. Compare this to, for instance, Van Dyke *et al*. (2022), who showed that rainfall changes largely affect the conditions for species coexistence. This work required a combination of theoretical approach and sophisticated field experiments involving six plant species and 106 planting plots subjected to two environmental treatments over two years. We thus expect this simplification provided by LV-map to generate many novel insights in the study of community dynamics. Our ability to directly query the *per capita* interaction strength and sign, which has been indicated to profoundly affect the niche differences between species in theoretical studies (Chesson, 2000; Saavedra *et al*., 2017; Song *et al*., 2020), such that greater deviations in species’ niches increase the possibility for coexistence. In addition, LV-map will also help confirm insights into the intrinsic growth rates, which are known to impact fitness differences, where smaller differences enable species coexistence (Chesson, 2000; Saavedra *et al*., 2017; Song *et al*., 2020).

Applying LV-map to experimental data, we were able to detect the allocative trade-offs between intrinsic growth rates and the *per capita* interaction strength, which essentially determines evolutionary outcomes in populations and communities. In fact, coexistence status can change as evolutionary processes direct within-species variations of intrinsic growth rates and *per capita* interaction coefficients (Lankau, 2011; Hart *et al*., 2019). Here, we show that fast-growing clones exhibit higher intraspecific competition and are more likely to be eaten by predators than slow-growing ones, i.e. there is a trade-off between growth versus competition and defence. We thus expect LV-map to provide deeper insights into underlying evolutionary processes.

While we did not use regularisation techniques for the LV-map in this study, appropriate techniques have been proposed, validated (Cenci *et al*., 2019), and can be applied if needed. Additionally, interpreting the inferred parameters requires caution, as unexpected results have been demonstrated by both the LV-map (Figure 4g) and S-map method (Deyle *et al*., 2016). In particular, in the occurrence of migration, the inferred *r*_*i*_(*t*) may no longer represent the intrinsic growth rate as it now encompasses both emigration and immigration. In the future, demographic characteristics of populations should also be considered in this model, as population dynamics may be structured in terms of age or sex, meaning that it may not be straightforward to interpret the parameters without consideration for these characteristics.

Overall, the LV-map is a promising approach for resolving many ecological and evolutionary questions while avoiding the time-consuming, labour-intensive, and disruptive isolation of organisms from their natural context. We expect that our proposed approach for inferring intrinsic growth rates and *per capita* interactions could pave the way for a broader understanding of ecological dynamics by allowing the use of time-series data from a range of natural and experimental communities. This feature alone should further improve our understanding of how species, phenotypes or genetic lineages coexist in complex ecosystems, and the mechanisms governing biodiversity. Given the increasing amount of time-series data being collected worldwide across systems (Benincà *et al*., 2015; Ehrlich & Gaedke, 2020; Merz *et al*., 2023), the broad applicability of this approach should help improve our overall understanding of the changing dynamics of ecosystems in our increasingly changing world.

## Acknowledgements

This work was funded by the Swiss National Science Foundation (Sinergia grant no CR-SII5 202290) to FP and RPR. RPR also acknowledges the Swiss National Science Foundation grant no 31003A 182386. We thank Ewa Merz, Marta Reyes, and Stefanie Merkli for preparing the high-frequency lake dataset. We are grateful to Serguei Saavedra, Luis J. Gilarranz, and Nicolas Loeuille for comments and discussion.

## Supporting Information for

### Standard error of the parameters

The standard error (SE) of the intercept (intrinsic growth rate; *r*_*i*_) and coefficient (inter and intraspecific interaction; *α*_*ij*_) follows the statistics of conventional multivariate regression method. The linear regression assumes that the deviation of the response variable 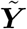 from its predicted values 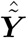 follows a normal distribution, therefore for each column *i* of the response matrix 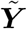 we have

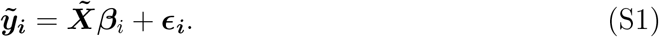

The vector ***β***_*i*_ corresponds to the column *i* of the matrix of parameters ***β***. The residuals ***ϵ***_***i***_ are assumed to be independent and identically distributed and follow a normal distribution with mean zero and variance 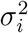. The residual variance 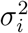 can be estimated by

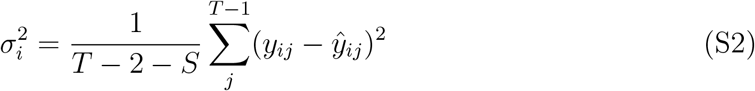

Then the estimation 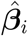 of the parameters ***β***_*i*_ follows a normal distribution of mean ***β*** and variance-covariance matrix given by

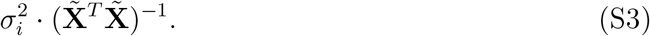

And thus, the square roots of the diagonal elements of this variance-covariance matrix are the standard error of 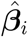 See chapter 3.2 of Hastie *et al*. (2001) and Chapter 3.3 of Mardia *et al*. (1979). The computation of the standard errors is done at each time point *t*.

### Cross validation

The weighting parameters, *θ*, is estimated using cross-validation technique. As the data points in time-series data are dependent, we cannot use classical cross validation methods that randomly split the training data (or in sample data) and the test data (or out of sample data). Instead, we use the cross validation on rolling origin forecast.

This cross validation technique uses the first-*t* observation of population abundance (**n**(*l*), for *l* = 1, … *t*) to predict the population abundance 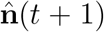 at time step *t* + 1. This process of prediction is iterated for *t* from *T*_*s*_ to *T −* 1. The initial time step *T*_*s*_ is usually chosen as 10% of the total number of time point *T*, i.e., *T*_*s*_ = round(*p · T*) with *p* = 0.1. The best *θ* is the one minimizing the root mean sum of error squares (RMSE). The expression of the RMSE is given by

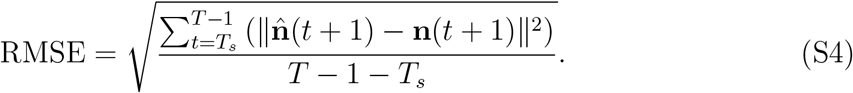

The prediction of 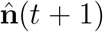 is done as follows. Knowing the abundances from time 1 to time *t*, we can estimate 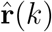 and 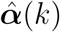 from time 1 to time *t −* 1. Note that we cannot estimate **r**(*t*) and ***α***(*t*), as it would require the knowledge of **n**(*t* + 1). Consequently, we use 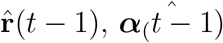 and **n**(*t*) to estimate 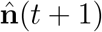 as follows

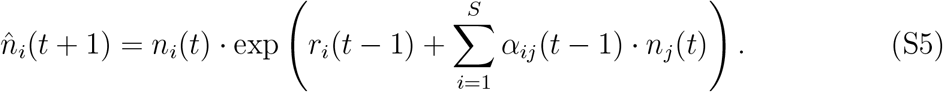

### Synthetic data

Synthetic data is simulated from a discrete time Lotka-Volterra model with three competing population and environmental noise. The equation is given by:

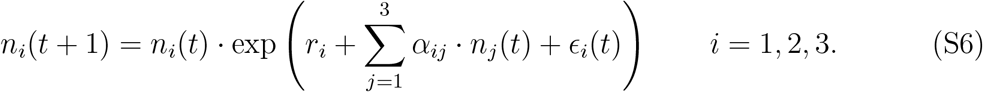

The parameters *r*_*i*_ and *α*_*i*_*j* are the intrinsic growth rate and the *per capita* interaction strengths, respectively. The random variables *ϵ*_*i*_(*t*) represent the environmental noise, which are drawn independently at random from a centred normal distribution and of standard deviation proportional to the *r*_*i*_, i.e., *ϵ*_*i*_(*t*) *∼ 𝒩* (0, *r*_*i*_ *· σ*). The parameter *σ* determine the overall environmental noise level. We use all points that are generated by the simulation.

### Experimental data

Time-series experimental data of Blasius *et al*. (2020) are obtained from open-source data, which the author share on FigShare https://doi.org/10.6084/m9.figshare.10045976.v1. We interpolate the missing data points. In particular, two missing points for experiment C2 and C3 (i.e. clone 2 and clone 3), and seven missing points for experiment C1 (i.e. clone 1). In experiment C1, we replace the zero density of rotifers by the minimum value of rotifer densities and divide it by 8. In addition, the experiment with algal clone 1 lasted for 350 days, but we only kept data from the first 190 days to match the length of the other two experiments (190 days for experiments with algal clone 2 and 181 days for algal clone 3).

Time-series experimental data of Yoshida *et al*. (2003) are obtained using PlotDigitizer app https://plotdigitizer.com/app. In particular, we took screen shorts of the figures from the paper, uploaded them on the PlotDigitizer website, and manually extracted the values of the data points.

### Observational data

Plankton high-frequency data were collected from Lake Greifensee, Switzerland, by a dual-magnification dark-field imaging microscope Merz *et al*. (2021). Pelagic plankton images in the size range between *∼* 10 *µ*m and *∼* 1 cm were collected at 3 m depth for 10 minutes every hour, and abundances (as regions of interest per second, ROI/s) were aggregated (summed) per day. For this study, we used data collected between March 2019 and December 2022. We classified taxa using a deep-learning classifier (Kyathanahally *et al*., 2021) (the code for the classification can be found in https://github.com/kspruthviraj/Plankiformer), and focused on five aggregated groups of phytoplankton: Cyanobacteria, Green algae, Diatoms, Golden algae, and Cryptohytes. We interpolate the missing data (50 points out of 1382 points), and replace the values that are absolute zero with the min value of the time-series of the corresponding group and divide them by 8. In this way, we represent the absolute zero values with extremely small values to enable the application of the LV-map.

## Supplementary figures

**FIGURE S1.**
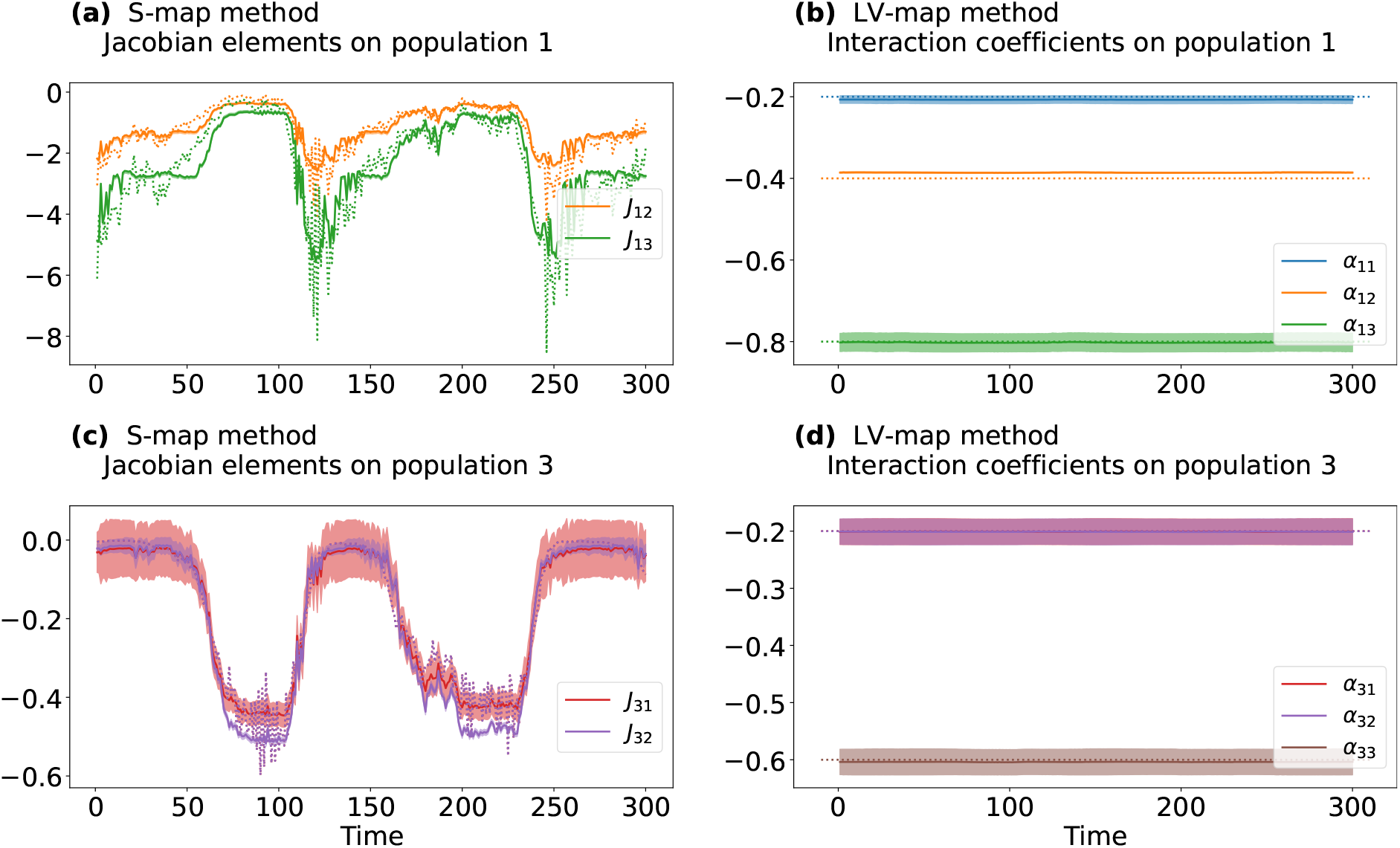
Estimation of parameters for population 2 and 3 of the cyclic Lotka-Volterra model. (a, c), Off-diagonal Jacobian elements for population 2 and 3. (b, d) *per capita* interactions for population 2 and 3. Solid lines with shaded areas represent estimated parameters and their standard errors, and dotted lines represent true values. Note that the *per capita* interactions of species 1 and 2 on species 3 are the same values, therefore the line red and purple lines overlap.

**FIGURE S2.**
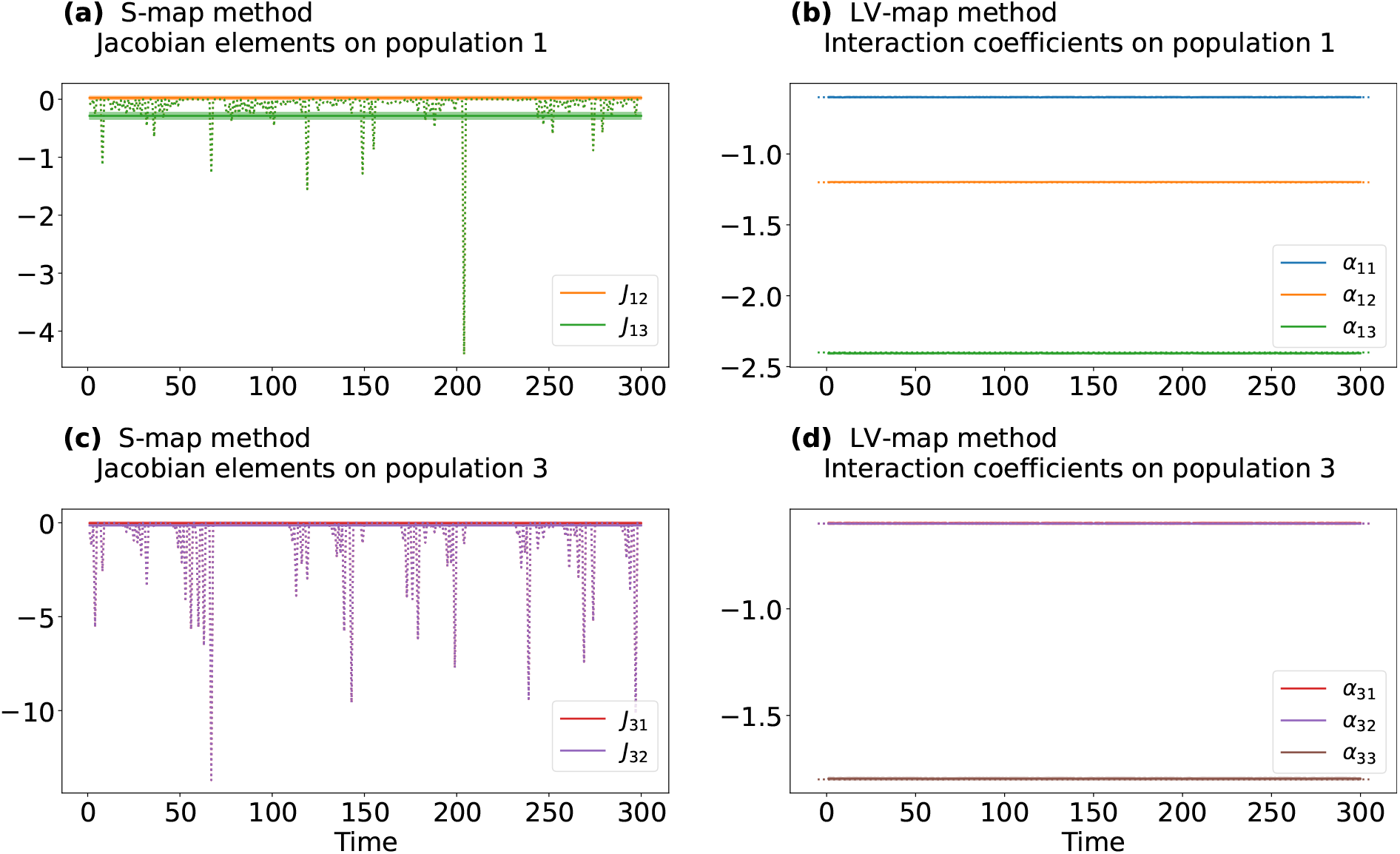
Estimation of parameters for population 2 and 3 of the chaotic Lotka-Volterra model. (a - c) Off-diagonal Jacobian elements for population 2. and 3. (b - d) *per capita* interactions for population 2 and 3. Annotations are similar to Fig S1.

**FIGURE S3.**
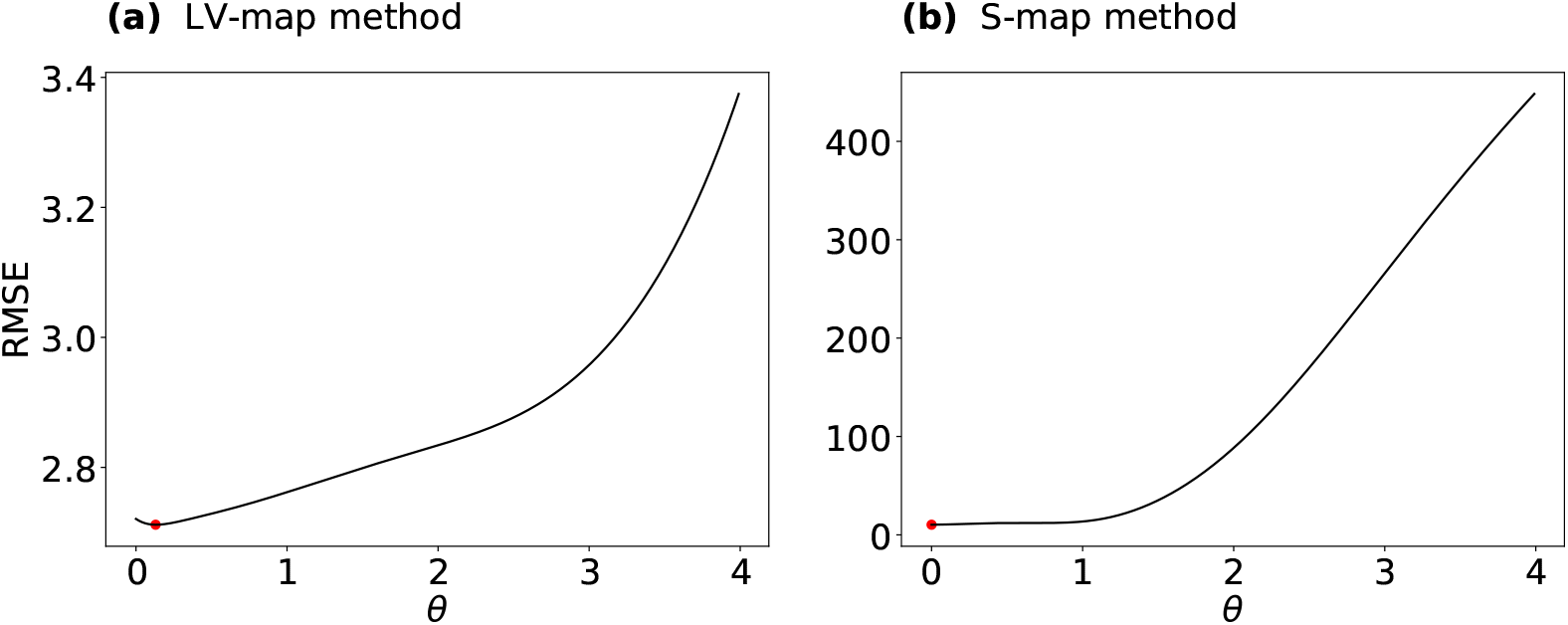
Cross validation results for the chaotic Lotka-Volterra model. (a), LV-map method. (b), S-map method. Red points indicate the value of *θ* with the smallest value of RMSE.

**FIGURE S4.**
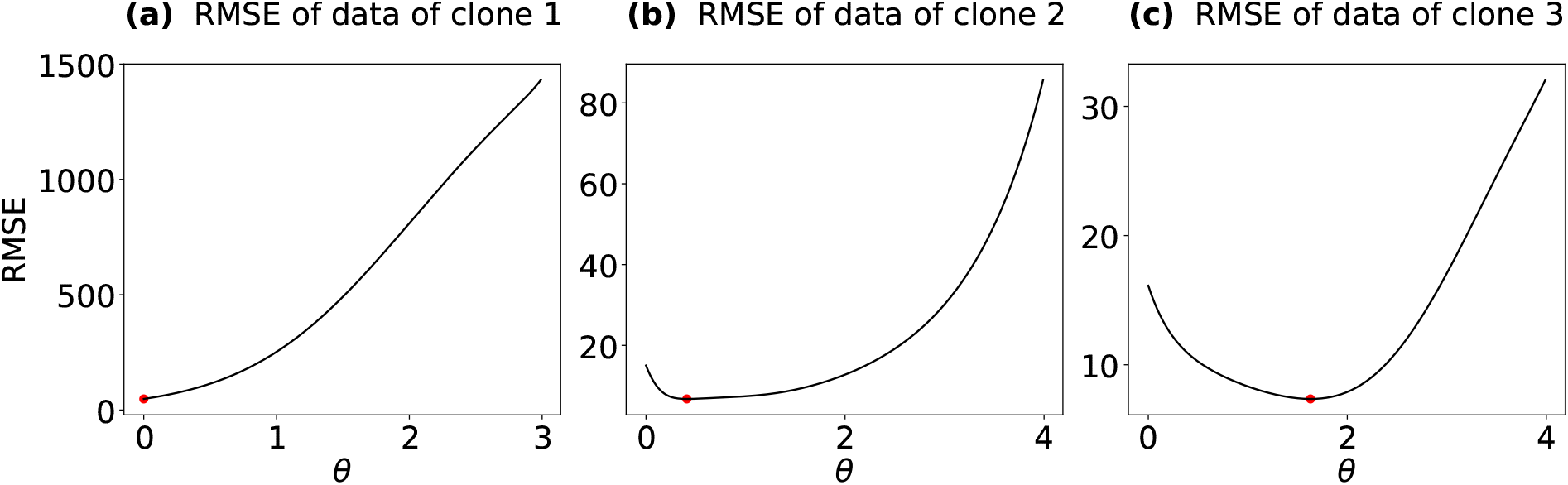
Cross validation results for experimental data of Yoshida *et al*. (2003). Results from experiment 1 (a), experiment 2 (b), and experiment 3 (c).

**FIGURE S5.**
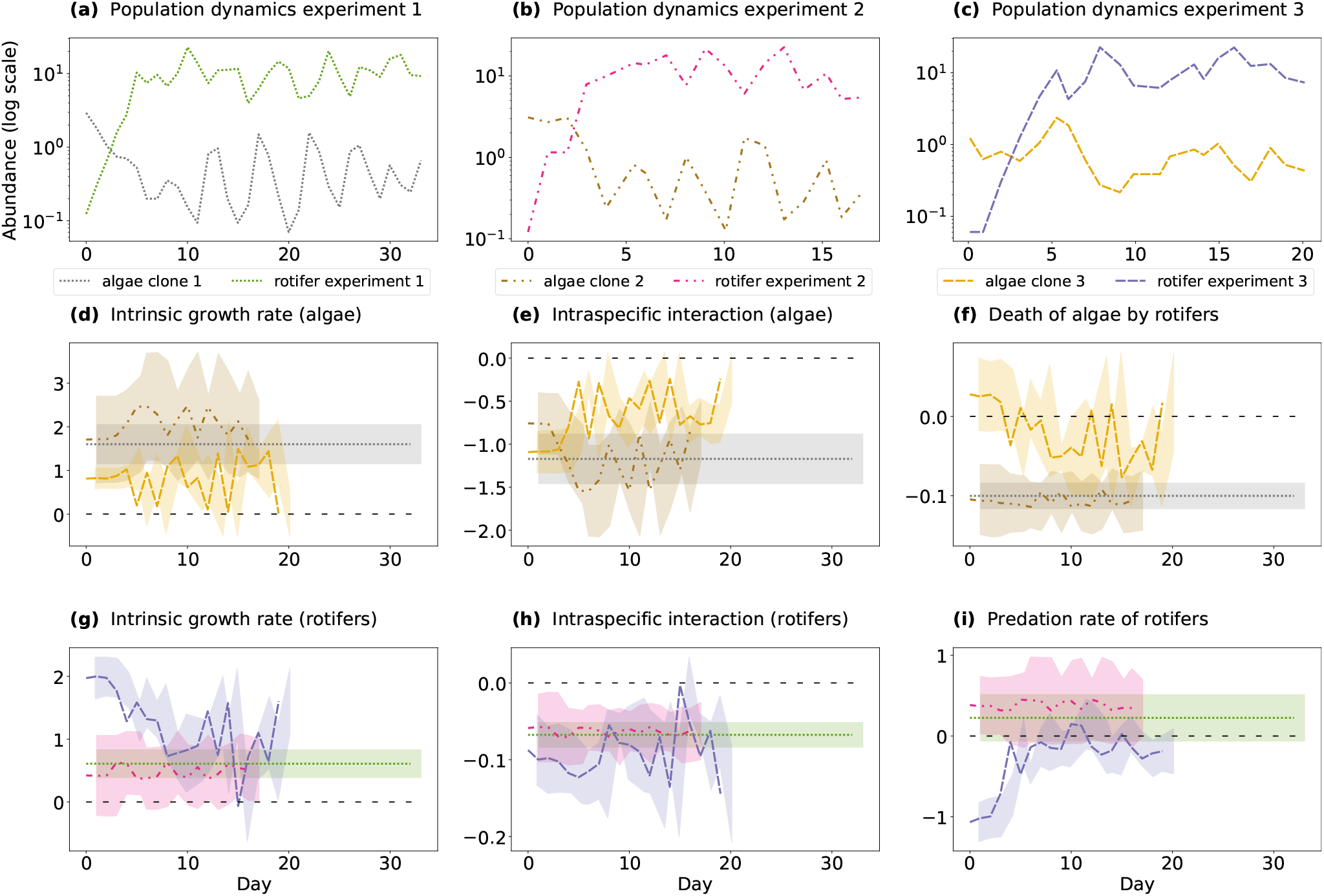
Time-series data of algae and rotifers from Yoshida *et al*. (2003). (a-c) Population dynamics of different clones. (d) Intrinsic growth rates of algae. (e), *per capita* intraspecific interactions between algae. (f), Effect of rotifers on algae. (g), Intrinsic growth rates of rotifers. (h), *per capita* intraspecific interactions between rotifers. (i), Effect of algae on rotifers. Different colours and line styles correspond to different clones (dotted for clone 1, dot-dot-dashed for clone 2, and dashed for clone 3). Shaded areas represent the standard errors.

**FIGURE S6.**
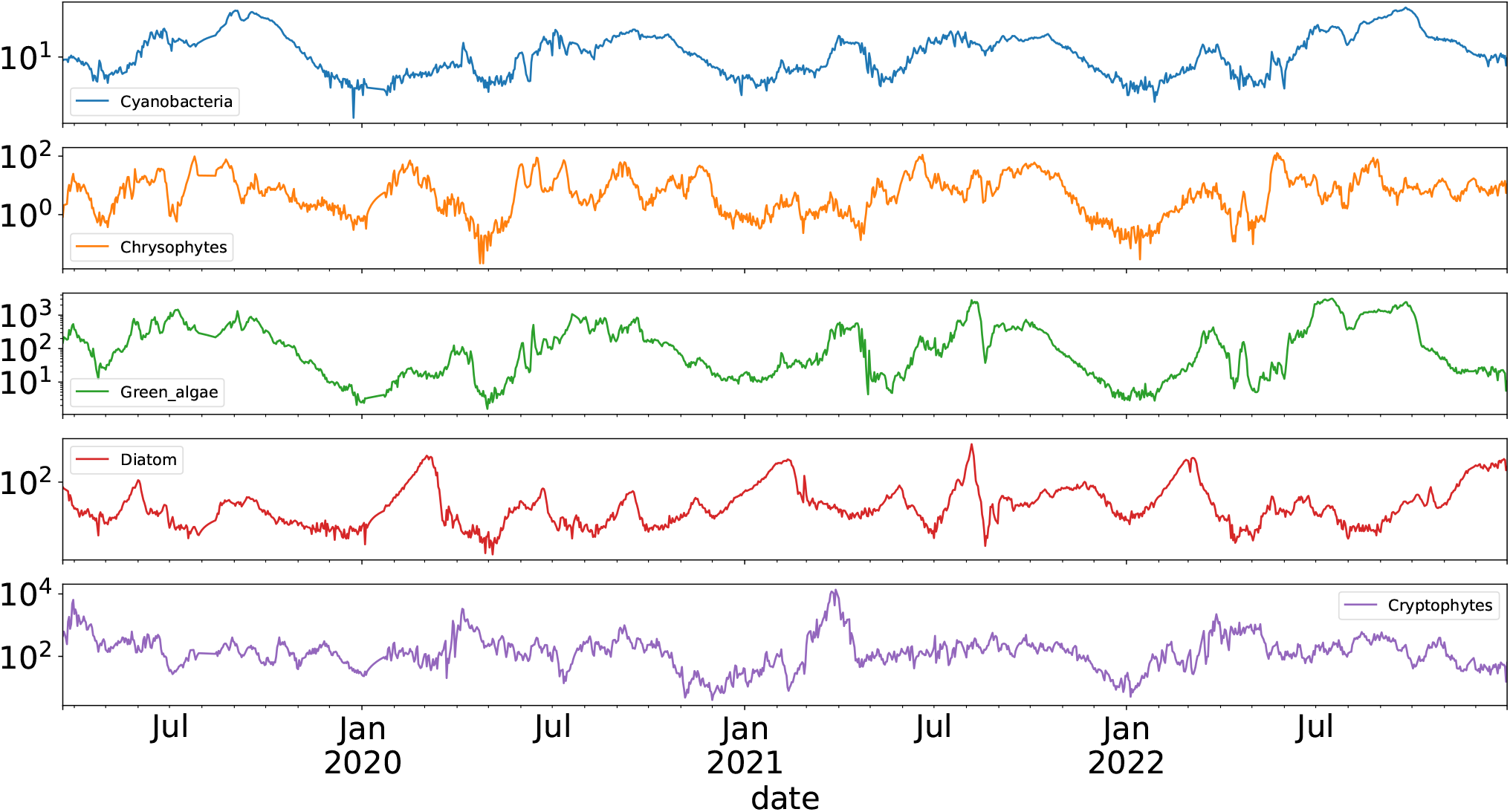
Time-series data of Cyanobacteria, Crysophyte, Green algae, Diatoms and Cryptophytes

**FIGURE S7.**
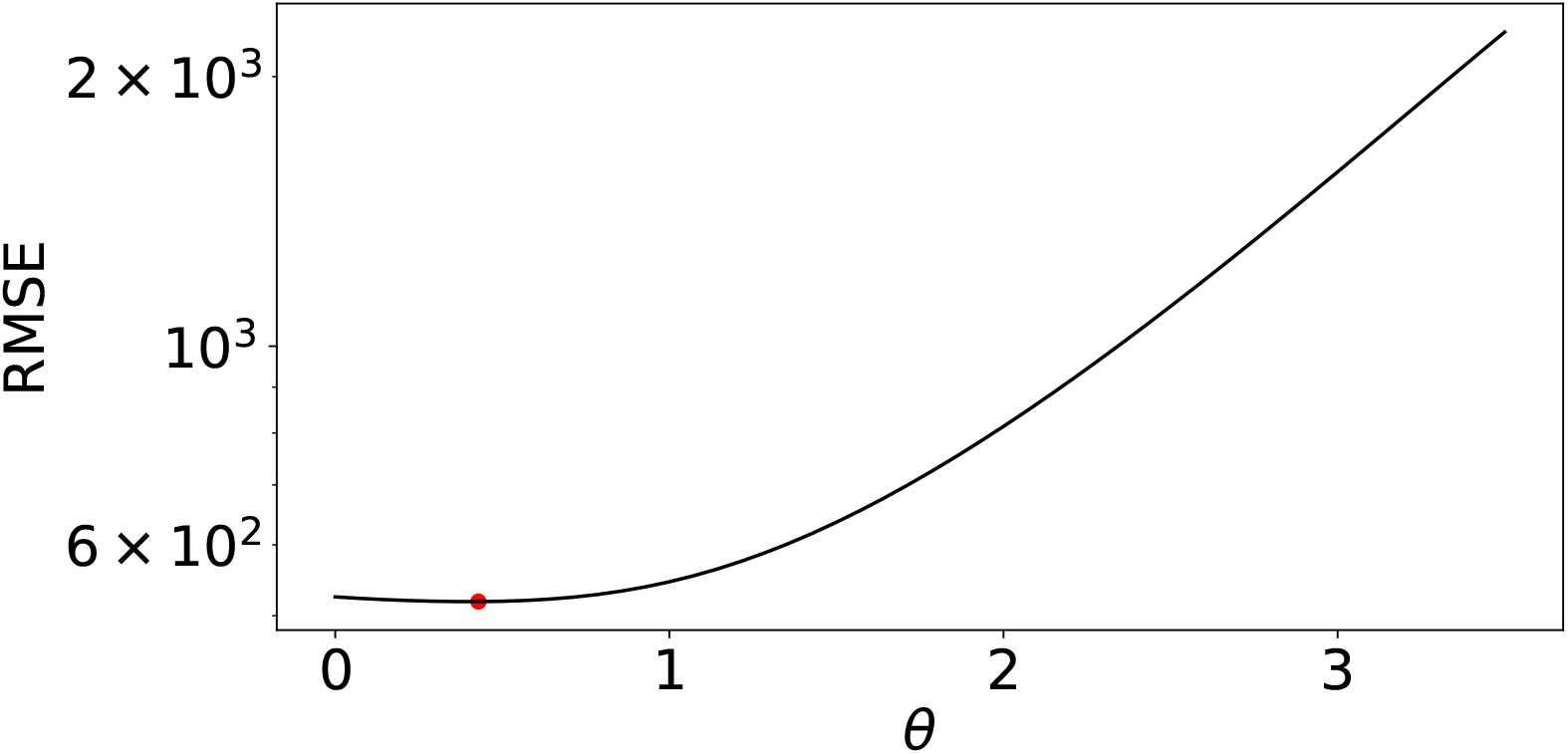
Cross validation results of the inference using lake data.

## References

Arditi, R., Tyutyunov, Y. V., Titova, L. I., Rohr, R. P. & Bersier, L.-F. (2021). The Dimensions and Units of the Population Interaction Coefficients. Frontiers in Ecology and Evolution, 9.

Baert, J. M., Eisenhauer, N., Janssen, C. R. & De Laender, F. (2018). Biodiversity effects on ecosystem functioning respond unimodally to environmental stress. Ecology Letters, 21, 1191–1199.

Baert, J. M., Janssen, C. R., Sabbe, K. & De Laender, F. (2016). Per capita interactions and stress tolerance drive stress-induced changes in biodiversity effects on ecosystem functions. Nat Commun, 7, 12486.

Bartomeus, I., Saavedra, S., Rohr, R. P. & Godoy, O. (2021). Experimental evidence of the importance of multitrophic structure for species persistence. Proceedings of the National Academy of Sciences, 118, e2023872118.

Bascompte, J. & Jordano, P. (2007). Plant-animal mutualistic networks: The architecture of biodiversity. Annual Review of Ecology, Evolution, and Systematics, 38, 567–593.

Bender, E. A., Case, T. J. & Gilpin, M. E. (1984). Perturbation Experiments in Community Ecology: Theory and Practice. Ecology, 65, 1–13.

Benincà, E., Ballantine, B., Ellner, S. P. & Huisman, J. (2015). Species fluctuations sustained by a cyclic succession at the edge of chaos. Proceedings of the National Academy of Sciences, 112, 6389–6394.

Berlow, E. L., Neutel, A.-M., Cohen, J. E., De Ruiter, P. C., Ebenman, B., Emmerson, M., Fox, J. W., Jansen, V. A. A., Iwan Jones, J., Kokkoris, G. D., Logofet, D. O., McKane, A. J., Montoya, J. M. & Petchey, O. (2004). Interaction strengths in food webs: issues and opportunities. Journal of Animal Ecology, 73, 585–598.

Blasius, B., Rudolf, L., Weithoff, G., Gaedke, U. & Fussmann, G. F. (2020). Long-term cyclic persistence in an experimental predator–prey system. Nature, 577, 226–230.

Cenci, S., Sugihara, G. & Saavedra, S. (2019). Regularized S-map for inference and forecasting with noisy ecological time series. Methods in Ecology and Evolution, 10, 650–660.

Chang, C.-W., Miki, T., Ushio, M., Ke, P.-J., Lu, H.-P.Shiah, F.-K. & Hsieh, C.-h. (2021). Reconstructing large interaction networks from empirical time series data. Ecology Letters, 24, 2763–2774.

Chesson, P. (2000). Mechanisms of Maintenance of Species Diversity. Annual Review of Ecology and Systematics, 31, 343–366.

Cohen, J. E., Schittler, D. N., Raffaelli, D. G. & Reuman, D. C. (2009). Food webs are more than the sum of their tritrophic parts. Proceedings of the National Academy of Sciences, 106, 22335–22340.

Cotter, S. C., Kruuk, L. E. B. & Wilson, K. (2004). Costs of resistance: genetic correlations and potential trade-offs in an insect immune system. Journal of Evolutionary Biology, 17, 421–429.

Deyle, E. R., May, R. M., Munch, S. B. & Sugihara, G. (2016). Tracking and forecasting ecosystem interactions in real time. Proceedings of the Royal Society B: Biological Sciences, 283, 20152258.

Ehrlich, E. & Gaedke, U. (2020). Coupled changes in traits and biomasses cascading through a tritrophic plankton food web. Limnology and Oceanography, 65, 2502–2514.

Godoy, O., Bartomeus, I., Rohr, R. P. & Saavedra, S. (2018). Towards the integration of niche and network theories. Trends in Ecology &amp; Evolution, 33, 287–300.

Hart, S. P., Turcotte, M. M. & Levine, J. M. (2019). Effects of rapid evolution on species coexistence. Proceedings of the National Academy of Sciences, 116, 2112–2117.

Hastie, T., Tibshirani, R. & Friedman, J. (2001). The Elements of Statistical Learning, chap. Linear methods for Regression. Springer New York Inc.

HilleRisLambers, J., Adler, P., Harpole, W., Levine, J. & Mayfield, M. (2012). Rethinking Community Assembly through the Lens of Coexistence Theory. Annual Review of Ecology, Evolution, and Systematics, 43, 227–248.

Holger Kantz, T. S. (2004). Nonlinear Time Series Analysis. 2nd edn. Cambridge University Press. ISBN 9780511078538; 0511078536; 9780521821506; 0521821509; 0521529026; 9780521529020.

Huisman, J., Codd, G. A., Paerl, H. W., Ibelings, B. W., Verspagen, J. M. H. & Visser, P. M. (2018). Cyanobacterial blooms. Nature Reviews Microbiology, 16, 471–483.

Kyathanahally, S. P., Hardeman, T., Merz, E., Bulas, T., Reyes, M., Isles, P., Pomati, F. & Baity-Jesi, M. (2021). Deep learning classification of lake zooplankton. Frontiers in Microbiology, 12.

Lankau, R. A. (2011). Rapid Evolutionary Change and the Coexistence of Species. Annual Review of Ecology, Evolution, and Systematics, 42, 335–354.

Laska, M. S. & Wootton, J. T. (1998). Theoretical Concepts and Empirical Approaches to Measuring Interaction Strength. Ecology, 79, 461–476.

Levine, J. M. & HilleRisLambers, J. (2009). The importance of niches for the maintenance of species diversity. Nature, 461, 254–257.

Lotka, A. J. (1925). Elements of physical biology. Williams & Wilkins.

Mardia, K., Kent, J. & Bibby, J. (1979). Multivariate analysis, chap. Normal distribution theory. Acad. Press.

Merz, E., Kozakiewicz, T., Reyes, M., Ebi, C., Isles, P., Baity-Jesi, M., Roberts, P., Jaffe, J. S., Dennis, S. R., Hardeman, T., Stevens, N., Lorimer, T. & Pomati, F. (2021). Underwater dual-magnification imaging for automated lake plankton monitoring. Water Research, 203, 117524.

Merz, E., Saberski, E., Gilarranz, L. J., Isles, P. D. F., Sugihara, G., Berger, C. & Pomati, F. (2023). Disruption of ecological networks in lakes by climate change and nutrient fluctuations. Nature Climate Change, 13, 389–396.

Paine, R. T. (1992). Food-web analysis through field measurement of per capita interaction strength. Nature, 355, 73–75.

Parain, E. C., Rohr, R. P., Gray, S. M. & Bersier, L.-F. (2019). Increased Temperature Disrupts the Biodiversity–Ecosystem Functioning Relationship. The American Naturalist, 193, 227–239.

Saavedra, S., Rohr, R. P., Bascompte, J., Godoy, O., Kraft, N. J. B. & Levine, J. M. (2017). A structural approach for understanding multispecies coexistence. Ecological Monographs, 87, 470–486.

Sibly, R. M. & Hone, J. (2002). Population growth rate and its determinants: an overview. Philos Trans R Soc Lond B Biol Sci, 357, 1153–1170.

Song, C., Rohr, R. P., Vasseur, D. & Saavedra, S. (2020). Disentangling the effects of external perturbations on coexistence and priority effects. Journal of Ecology, 108, 1677–1689.

Strauss, S. Y., Rudgers, J. A., Lau, J. A. & Irwin, R. E. (2002). Direct and ecological costs of resistance to herbivory. Trends in Ecology & Evolution, 17, 278–285.

Sugihara, G. (1994). Nonlinear Forecasting for the Classification of Natural Time Series. Philosophical Transactions: Physical Sciences and Engineering, 348, 477–495.

Turchin, P. (1999). Population Regulation: A Synthetic View. Oikos, 84, 153–159.

Van Dyke, M. N., Levine, J. M. & Kraft, N. J. B. (2022). Small rainfall changes drive substantial changes in plant coexistence. Nature, 611, 507–511.

Vandermeer, J. H. (1969). The Competitive Structure of Communities: An Experimental Approach with Protozoa. Ecology, 50, 362–371.

Vincent, T. L. & Brown, J. S. (2005). Evolutionary Game Theory, Natural Selection, and Darwinian Dynamics. Cambridge University Press, Cambridge. ISBN 978-0-521-84170-2.

Volterra, V. (1931). Lecon sur la théorie mathematique de la lutte pur la vie. Gauthier Villars.

Yoshida, T., Hairston, Jr, N. G. & Ellner, S. P. (2004). Evolutionary trade-off between defence against grazing and competitive ability in a simple unicellular alga, chlorella vulgaris. Proc. Biol. Sci., 271, 1947–1953.

Yoshida, T., Jones, L. E., Ellner, S. P., Fussmann, G. F. & Hairston, N. G. (2003). Rapid evolution drives ecological dynamics in a predator–prey system. Nature, 424, 303–306.

